# Better long-term learning ability is predicted by higher surface folding of the human premotor cortex

**DOI:** 10.1101/2023.08.18.553797

**Authors:** Marco Taubert, Gabriel Ziegler, Nico Lehmann

**Affiliations:** Department of Sport Science, Institute III, Faculty of Humanities, Otto von Guericke University, Zschokkestraße 32, 39104 Magdeburg, Germany; Department of Neurology, Max Planck Institute for Human Cognitive and Brain Sciences, Stephanstraße 1a, 04103 Leipzig, Germany; Germany German Center for Neurodegenerative Diseases (DZNE), Leipziger Straße 44, 39120 Magdeburg, Germany; Center for Behavioral and Brain Science (CBBS), Otto von Guericke University, Universitätsplatz 2, 39106 Magdeburg, Germany; Institute of Cognitive Neurology and Dementia Research, Otto von Guericke University, Leipziger Str. 44, 39120 Magdeburg, Germany; Collaborative Research Center 1436 Neural Resources of Cognition, Otto von Guericke University, Leipziger Str. 44, 39120 Magdeburg, Germany

## Abstract

The capacity to learn enabled the human species to adapt to various challenging environmental conditions and pass important achievements on to the next generation. A growing body of research suggests links between neocortical folding and numerous aspects of human behaviour, but their impact on enhanced human learning capacity remains unexplored. Here we leverage multiple training cohorts to demonstrate that higher levels of premotor cortical folding reliably predict individual long-term learning gains in a challenging new motor task, above and beyond initial performance differences. Individual folding-related predisposition to motor learning was found to be independent of cortical thickness and several intracortical microstructural parameters, but dependent on larger cortical surface area. We further show that learning-relevant features of cortical folding occurred in close spatial proximity to practice-induced structural plasticity and were primarily localized in hominoid-specific frontal tertiary sulci. Our results suggest a new link between neocortical surface folding and human behavioural adaptability.

## Introduction

Cortical folding is a highly complex developmental process that depends on the genotype^1^ and reflects the functional organization of the cortex ^2–6^, with striking similarities but also numerous differences between individuals and across species ^7,8^. It has been suggested that cortical folding evolved to fit a larger sheet-like cortex into a compact cranial space and to keep cortical nerve fiber connections short ^9–11^. This evolutionary expansion and folding of the human neocortex, especially in associative cortices, likely enhanced the neurocomputational capacities required for complex social interaction, tool-making and mobility ^12^.

The impact of cortical folding on behaviour has fascinated early neuroanatomists ^13–15^ and stimulates contemporary research in diverse fields such as biology, anthropology or cognitive neuroscience ^12,16–18^. The dominant view is that higher levels of cortical folding are directly linked to improved cognitive performance both within and across species ^11,14,19,20^. The number and interconnectivity of horizontally arranged cortical columns limit the information processing capacities of neural networks and its potential power for high cognitive performance ^9^. Patients with certain neurodevelopmental disorders present cortical folding abnormalities and cognitive deficits ^21^ and cross-sectional studies in healthy populations demonstrate positive correlations between normative cortical morphology and behavioural performance (most frequently with parameters of ‘intelligence’) but with varying small to moderate effect sizes ^19,22–25^. However, evidence for associations between cortical folding and longitudinal trajectories of behavioural change is still missing. We here exploit multi-cohort longitudinal data to test the hypothesis that cortical folding in the motor system might form a potential predisposition for intra-individual performance gains during motor practice.

A high level of behavioural performance might result from individual brain and body development, task-specific practice and/or previous experiences with similar tasks. The capability to improve performance through practice enabled the human species to adapt to various challenging environmental conditions and pass important achievements on to the next generation ^26,27^. It has been hypothesized that high human performance does not directly result from evolved brain features alone, but rather from an interaction between fertile learning environments (with rich opportunities for self-regulated and socially mediated learning) and remarkable learning capacities provided by the brain ^28,29^. Motor learning induces brain plasticity^30^ but behavioural genetics research also suggests that practice increases the relative importance of genetic influences on performance and reduces the effects of environmental variation resulting from different prior experiences ^31,32^. Therefore, learning in the human brain appears to be mediated by certain predispositions and practice-induced neural plasticity in the cortical and subcortical gray and white matter ^22,25,33,34^. However, no study to date investigated whether neocortical folding relates to motor learning capability. Building on recent developmental studies of behaviourally relevant features of cortical shape ^5^, genetics research on motor learning^31^ and our own work on motor learning-induced cortical plasticity ^35^, we hypothesize that individual variations in cortical folding does predict the individual potential to learn a new motor task and that such folding variations colocalize with learning-induced neural plasticity.

In the human brain, local geometric features of the cortical surface appear to fundamentally constrain differences in brain function ^36^. Cortical geometric features, such as local cortical curvature, can be assessed *in-vivo* using magnetic resonance imaging (MRI). Curvature-based metrics were used in previous cross-sectional studies to relate the local folding properties of cerebral regions to human behavioural performance ^37,38^ and to individual genotype ^39^. Moreover, surface-based metrics of cortical folding, such as local gyrification index, are particularly sensitive to differences in the size of the cortical surface buried within sulci ^40,41^. Morphometric analyses of cortical sulci in associative brain regions recently revealed a new role of hominoid-specific tertiary sulcus morphology for cognitive performance ^8,41–44^. Here we adopt a multi-scale approach to test the impact of local cortical folding on motor learning in multiple samples. In cortical regions with learning-relevant geometrical features (cortical curvature), we further investigate the contribution of cortical surface area, cortical thickness and intracortical microstructure (assessed using myelin-sensitive magnetization transfer saturation and neurite density index) to cortical geometry as well as the morphometric properties of closely overlapping tertiary sulci.

Specifically, we test for aptitude-treatment interactions ^45^ to disentangle the contributions of cortical folding either to superior (absolute) performance or adaptive capability (performance gain). The joint analysis of multimodal MRI data from three separate motor learning experiments ^35,46–48^ allows us to examine individual differences in motor learning in a challenging balance task over a practice period of 4 to 6 weeks ^49^ (Fig. 1A). We hypothesize that a contribution of cortical folding to superior performance would manifest in positive correlations with absolute performance differences while a contribution to superior learning capability would manifest in positive correlations with intraindividual performance gains (above and beyond initial performance differences).

**Fig. 1.**
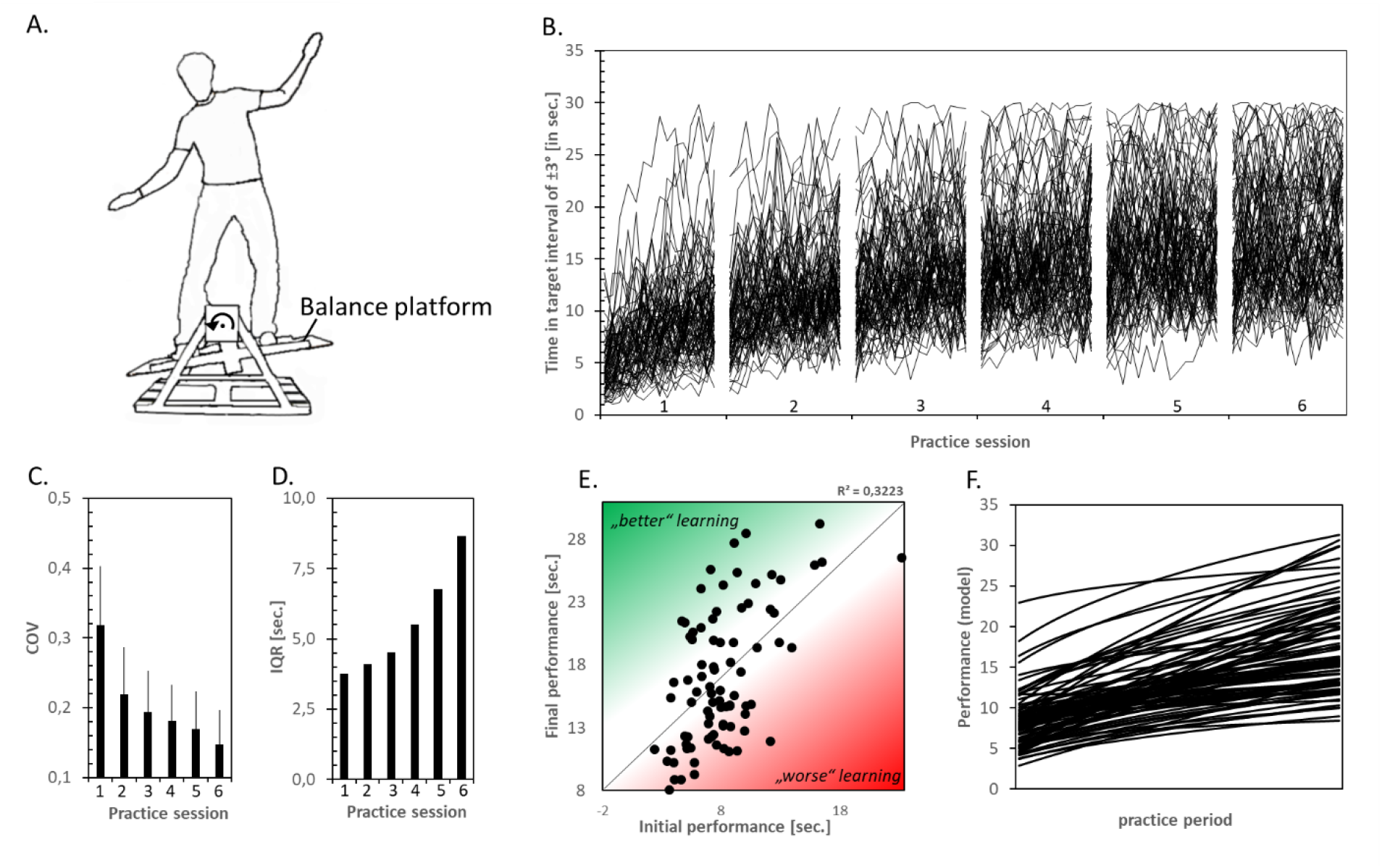
Behavioural data. Motor learning task, performance improvements, performance stabilization and increased inter-individual differences in motor learning over 6 practice sessions (*N*=84, mean age 24.6 years, age range 19-35 years, 57 women, mean height 174 cm, height range 153-191 cm, all participants were right-handed). (A) We tested motor learning of a challenging postural task. Participants were instructed to keep a seesaw-like moving stabilometer balance platform in a horizontal target interval (±3°) as long as possible during a trial length of 30 s. (B) Motor performance was measured as the time (in seconds) in which participants kept the board within the target interval in each of 15 practice trials per session (see Supplemental Video files for motor performance of participants at the beginning and end of practice). (C) Decrease in trial-to-trial variability (coefficient of variation, COV) of session-specific motor performance. (D) Increase of the interquartile range (IQR) of session-specific between-person variation in motor performance. IQR increased from 3.7 seconds at session 1 to 8.7 seconds at session 6. (E) From the first to the sixth session, participants tended to maintain their performance rank (correlation between initial and final performance, *R*^*2*^ = 0.322, *p* < .001) but there were large individual learning differences in learning (green/red: higher/lower performance than predicted from baseline). (F) Modeled individual learning curves over sessions using parameters of the power function (see main text).

## Results

### Long-term motor learning improves performance, reduces intra-individual performance variability and enhances inter-individual performance differences

Participants learned a whole-body balance task in six practice sessions spread over four to six weeks (Fig. 1A, B). Throughout the practice period motor performance increased continuously (main effect of session *F*(5, 415) = 202.61, *p* < .001, *ηp*^*2*^ = 0.709) with significant performance gains across the six practice sessions (all post-hoc comparisons between time points were significant at *p* < .001, Bonferroni corrected for multiple comparisons). Intraindividual (trial-to-trial) variability decreased (main effect of session *F*(5, 415) = 109.89, *p* < .001, *ηp*^*2*^ = 0.570, Fig. 1C) and absolute between-person performance differences (IQR) increased during practice (Fig. 1D). We found considerable inter-individual variability in motor learning (Fig. 1E). To relate variations in cortical folding to differences in the rate of motor learning, we first fitted a general power function

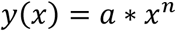

to the session-specific mean performance scores of each participant (Fig. 1F). The intercept *a* of the power function represents initial performance, while the exponent *n* reflects the individual learning rate and *x* is time. The general power function yielded an adequate fit to the individual learning data with a median coefficient of determination of *R*^2^ =.90. In accordance with the literature 50, initial performance *a* negatively predicted learning rate *n* (*R*^*2*^ = 0.350, *p* < .001, Fig. S1). We therefore adjusted learning rate *n* for inter-individual differences in initial performance *a* ^51^. We further use term ‘learning rate’ for this during all subsequent analyses.

### Cortical folding predicts inter-individual differences in long-term motor learning

We quantified vertex-wise cortical curvature to measure local cortical folding ^52^. Larger values indicate higher degrees of local cortical curvature. We then tested for correlations between higher cortical curvature and steeper learning curve (learning rate *n* adjusted for initial differences *a*), superior initial performance (intercept *a*), higher short-term adaptations during session 1 and higher asymptotic performance in session 6. All analyses were adjusted for age, gender, body height, study, and total intracranial volume (see covariate correlation matrix in Fig. S2).

We did not observe significant correlations between local curvature and initial performance or short-term adaptations (Figs. S3, S4). Instead, a steeper learning rate *n* was positively associated with higher cortical curvature in the left pre-supplementary/supplementary motor area (pre-SMA/SMA, peak at x=-13, y=18, z=63, T=5.97, FWE correction at p < .05, nonparametric t-statistic with 5000 permutations, see Figs. 2A,B and S5). The moderate effect size was consistent across the three sub-samples (Fig 2C). These positive (sample and subsample) correlations were reproduced in a second MRI scan of the same participants (Fig. S6). These subsequent analyses revealed that approximately 30% of the variance in adjusted learning rates was explained by differences in cortical curvature in pre-SMA/SMA (*R*^*2*^ = 0.30, *p* < .001, N = 84). The positive correlation between curvature and performance or gain increased during practice (Fig. 2E). In addition, cortical curvature consistently predicted learning rates within demographic, anthropometric, and performance-specific subcategories of the dataset (Figs. S7 and S8). Asymptotic (final) performance showed a non-significant trend for an association with cortical curvature in left pre-SMA/SMA (local maximum at x=-15, y=20, z=62, T=4.40, FWE-corrected p = .053) and a significant association in a small cluster in left supramarginal gyrus (local maximum at x=-59, y=-56, z=21, T=4.55, FWE correction at p < .05, see Fig. S9). In order to confirm the links between cortical folding, learning rates and final performance, we used structural equation modeling (see Materials for SEM fit indices) to show that the effect of cortical folding on final performance was mediated via learning rate *n* (Fig. 2D).

**Fig. 2.**
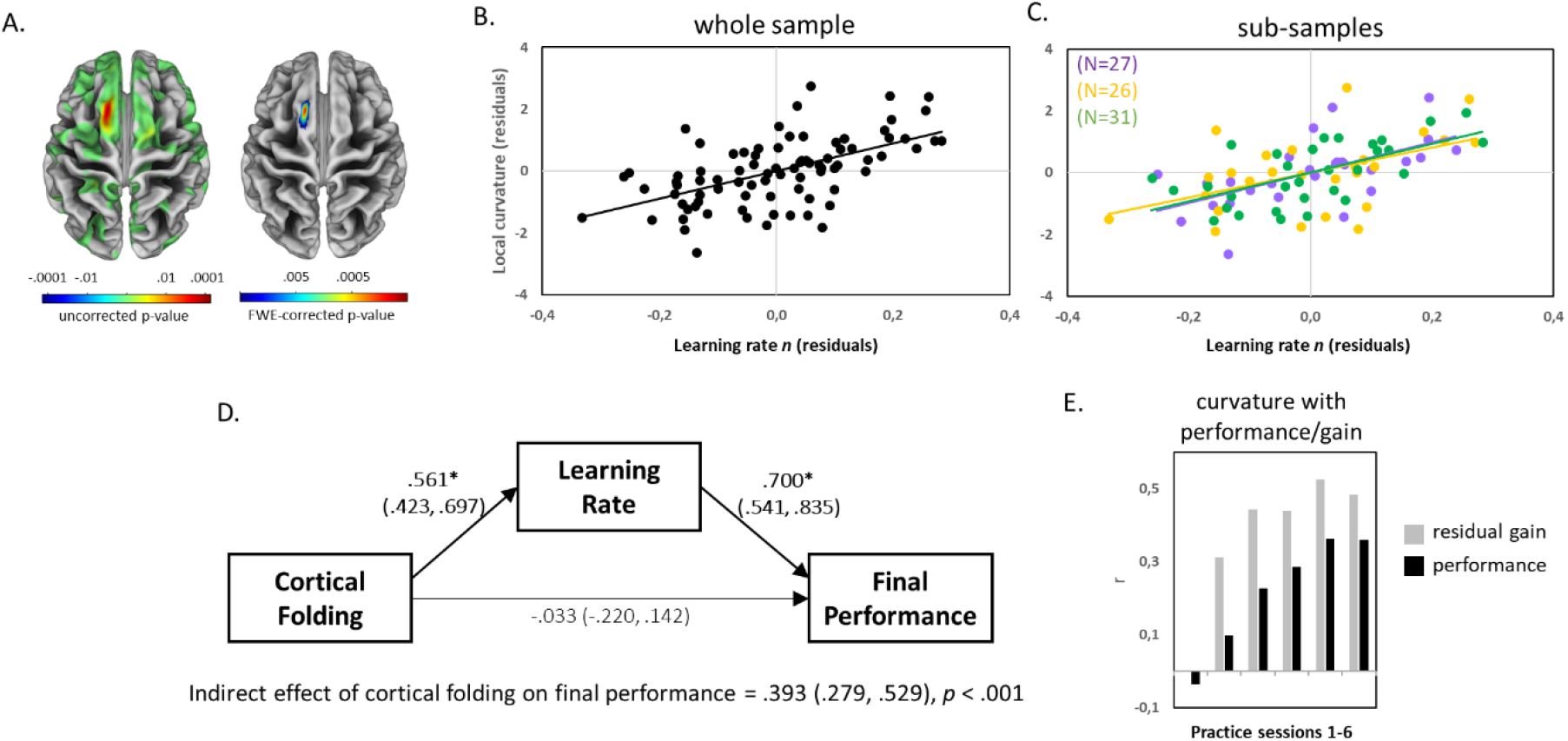
Cortical folding predicts learning. Results of whole-brain correlation of vertex-wise cortical curvature and learning rate. (A) Uncorrected results at *p* < .001 (left) and family-wise error-corrected results at *p* < .05 (right) were projected onto a template brain showing variations in sulcus depth. (B) Positive correlation of residual cortical folding (in the cluster representing the FWE-corrected effect in the exploratory analysis [A]) and learning rate. (C) Subsample results in the three independent learning experiments. (D) Structural equation model depicting relationships between cortical folding in pre-SMA/SMA (cluster from 2A, unadjusted for *a*), learning rate (adjusted for *a*) and final performance on session 6 (unadjusted for *a*). Standardized coefficients with 95% bootstrapped confidence intervals (CI) are represented on paths. (E) Pearson correlation coefficients between residualized cortical folding and motor performance (N = 84). Grey bars represent session-specific performance controlled for initial performance in session 1 (i.e., residual gain) and black bars represent correlations with actual session-specific performance. * indicate significant paths at *p* < .05 (with CIs not including zero).

### Individual folding-related predisposition to motor learning was independent of cortical thickness, but dependent on cortical surface area

At the macroscopic level, cortical folding depends on the size and thickness of the cortical sheet (surface area and cortical thickness, see ^53^). Thus, we tested the potential contributions of cortical surface area and cortical thickness to the observed relationship between cortical folding and learning rate using structural equation modeling (SEM).

Modelling results are shown in Figure 3B (see Materials for model fit indices). Within a larger region encompassing left pre-SMA/SMA (see Methods for ROI description), cortical surface area, but not cortical thickness, exerted an indirect effect on learning rate *n* via folding (indirect effect of surface area on *n*: 0.54 [95% CI = .305, .749], *p* < .001; no indirect effect of thickness on *n*: 0.02 [95% CI = -.076, .134], *p* = .686). In the context of this model, there was a direct effect of cortical folding on learning rate *n* (*R*^*2*^ = 0.21). Figure 3C shows a simple Pearson correlation between cortical folding and *n* (*R*^*2*^ = 0.16, *p* < .001). Importantly, the positive relationship between cortical folding and learning rate *n* remained significant when adjusting for differences in surface area and cortical thickness in a partial correlation analysis (*R*^*2*^ = 0.17, *p* < .001, Fig. 3D). In order to validate the effect of premotor cortical curvature on learning rate, we used a surface area-dependent gyrification index^40^ and found a spatial pattern of positive correlation in the same cortical region (Fig. S10). Thus, while surface area affected cortical folding, surface area-independent contributions to local cortical geometry also affected learning rate.

**Fig. 3.**
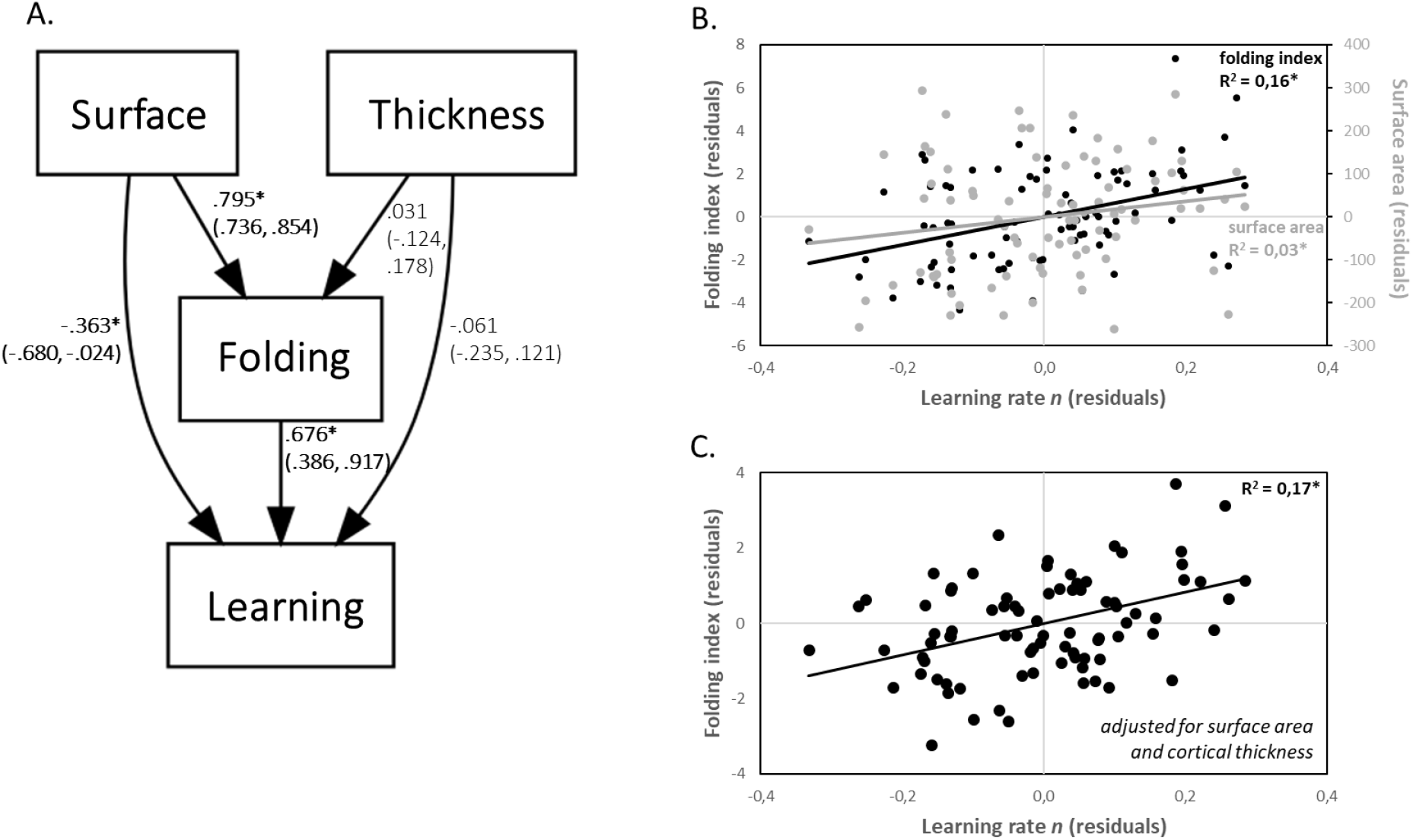
Cortical surface area, but not cortical thickness, is related to the effect of cortical folding on learning. Interrelationship between folding, thickness and surface area. (A) SEM model depicting the relationships between cortical folding (‘folding’), cortical surface area (‘surface’), cortical thickness (‘thickness’), and learning rate *n* (‘learning’) in the left caudal superior frontal gyrus. Standardized coefficients with 95% bootstrapped CIs are represented on paths. (B and C) Correlations between folding index and surface area with learning rate *n*. Folding index is either adjusted (C) or unadjusted (B) for differences in surface area and cortical thickness. Note that all variables used in the model and for correlation analyses were corrected for differences in age, gender, height, study, head coil, baseline performance, and total intracranial volume. * indicate significant paths/correlations at *p* < .05 (with CIs not including zero).

### Cortical folding ties to learning rates independent of cortical myelination and cortical neurite density

Cross-species comparisons do suggest that highly convoluted cortices have lower neuronal densities than less convoluted cortices ^54^. Also, the folding process in regions developing late during gestation (secondary and tertiary sulci) is likely to be mediated by intracortical microstructure ^6^ and biomechanical constraints ^55^. Intracortical myelination of deep cortical gray matter (GM), as measured by myelin-sensitive magnetization transfer saturation (MT), is a signature of cortical maturation in late adolescence and early adulthood ^56,57^. In order to test whether the folding effect on learning rate is significantly influenced by interindividual differences in intracortical microstructure, we measured myelin-sensitive MT saturation in superficial cortical to cortex-adjacent white matter compartments and intracortical neurite density index (NDI) of pre-SMA/SMA (*N* = 26; mean age 22.1 years, range 19-29 years, Fig. 4C). In line with previous studies, we observed a positive correlation between MT, in particularly in deep cortical GM, and chronological age in vertex-wise (Fig. S11) and ROI-wise correlation analyses (*R*^*2*^ = 0.33, *p* = .002, Fig. 4D). Importantly, we found no significant correlation between MT and learning rate *n* either using mean ROI (*R*^*2*^ ranged from 0.017 to 0.034, *p* > 0.36, Fig. 4E) or vertex-wise analyses (Fig. S9). Variations in MT had no impact on the association between cortical folding and learning rate *n* (partial *R*^*2*^ ranged from 0.26 to 0.27, all *p* < .009, Fig. 4F). In line with the MT analysis, learning rate *n* was not related to NDI values in pre-SMA/SMA. These results via imaging proxies indicate that the effect of higher cortical folding on steeper learning curves is less likely to be mediated by lower intracortical myelin content or neurite density across individuals.

**Fig. 4.**
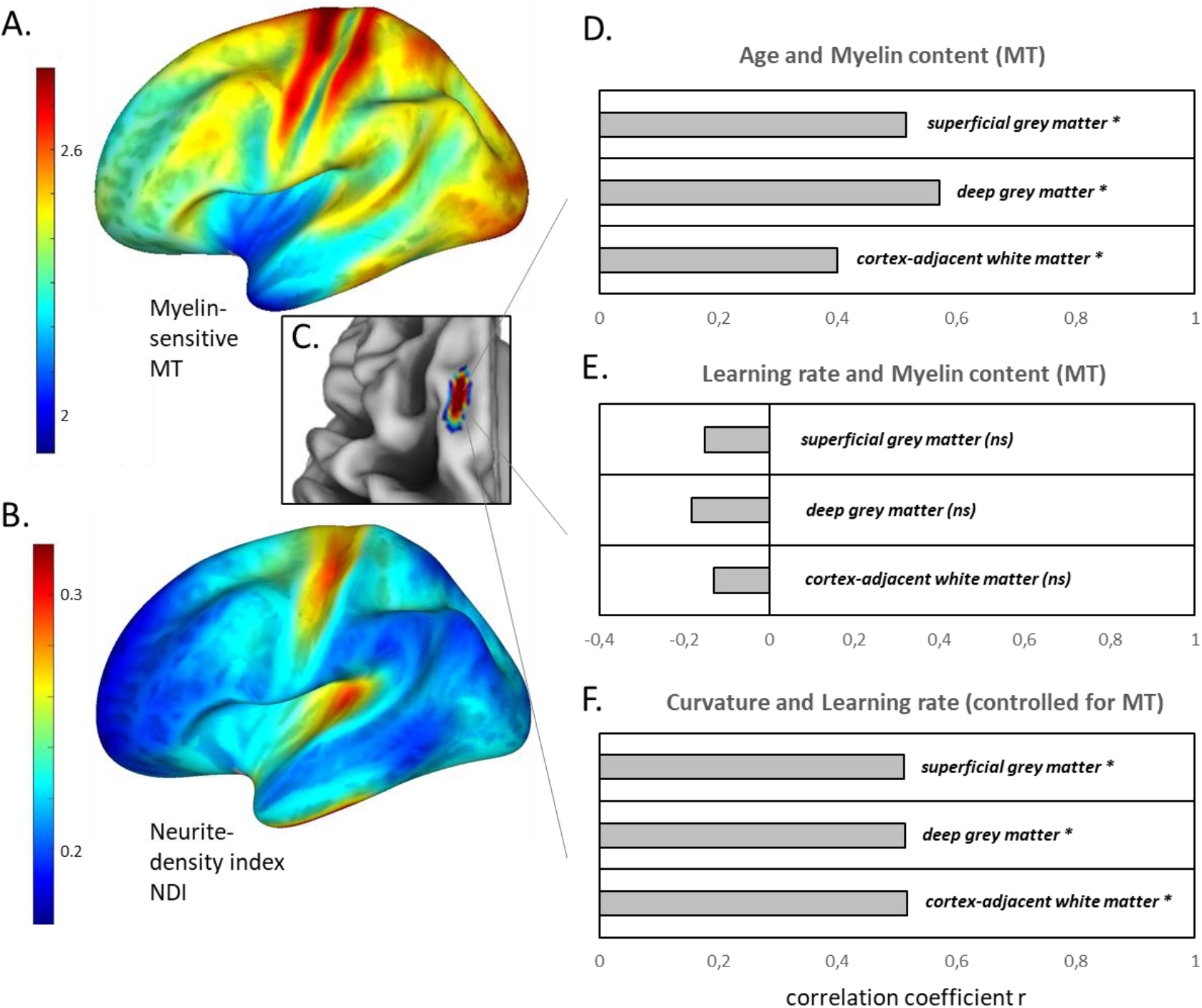
Cortical folding ties to learning rates independent of cortical myelination and cortical neurite density. Analysis of microstructural tissue properties of the premotor cortex. (A and B) Distribution of myelin-sensitive magnetization transfer saturation (MT) values (A) and the neurite density index NDI (B) across the left hemisphere. Color bars show regions of high MT or NDI in red (e.g., primary motor and somatosensory cortices) and regions of lower MT and NDI in blue (e.g., anterior prefrontal regions). Note the MT product-sequence-specific representation of MT values with a factor of 2. (C) MT and NDI values were analyzed in pre-SMA/SMA, the cluster in which cortical folding positively correlated with learning rate *n* (Fig. 2A). (D) Pearson correlations between MT in superficial GM, deep GM, and cortex-adjacent white matter with chronological age. (E) Pearson correlations between MT in superficial GM, deep GM, and cortex-adjacent white matter with learning rate *n*. (F) Partial correlations between cortical folding and learning rate adjusted for MT in superficial GM, deep GM, and cortex-adjacent white matter. * indicate significant correlations at *p* < .05, while ns indicates no significant correlation.

### Coincident effects of cortical folding and practice-induced plasticity

Our previous study showed structural gray matter plasticity in the pre-SMA/SMA after practice of the very same balance task ^35^ (Fig. 5A left). This gives us the opportunity to test the spatial coincidence of folding predispositions for learning and short-term learning-induced plasticity. Within the clusters that showed gray matter increases across the whole motor practice period (Fig. 2 in ^35^), higher cortical curvature in pre-SMA/SMA significantly predicted higher learning rate (peak at x=-15, y=18, z=59, T=5.64, FWE corrected *p*-value = .001, Fig. 5A right).

**Fig. 5.**
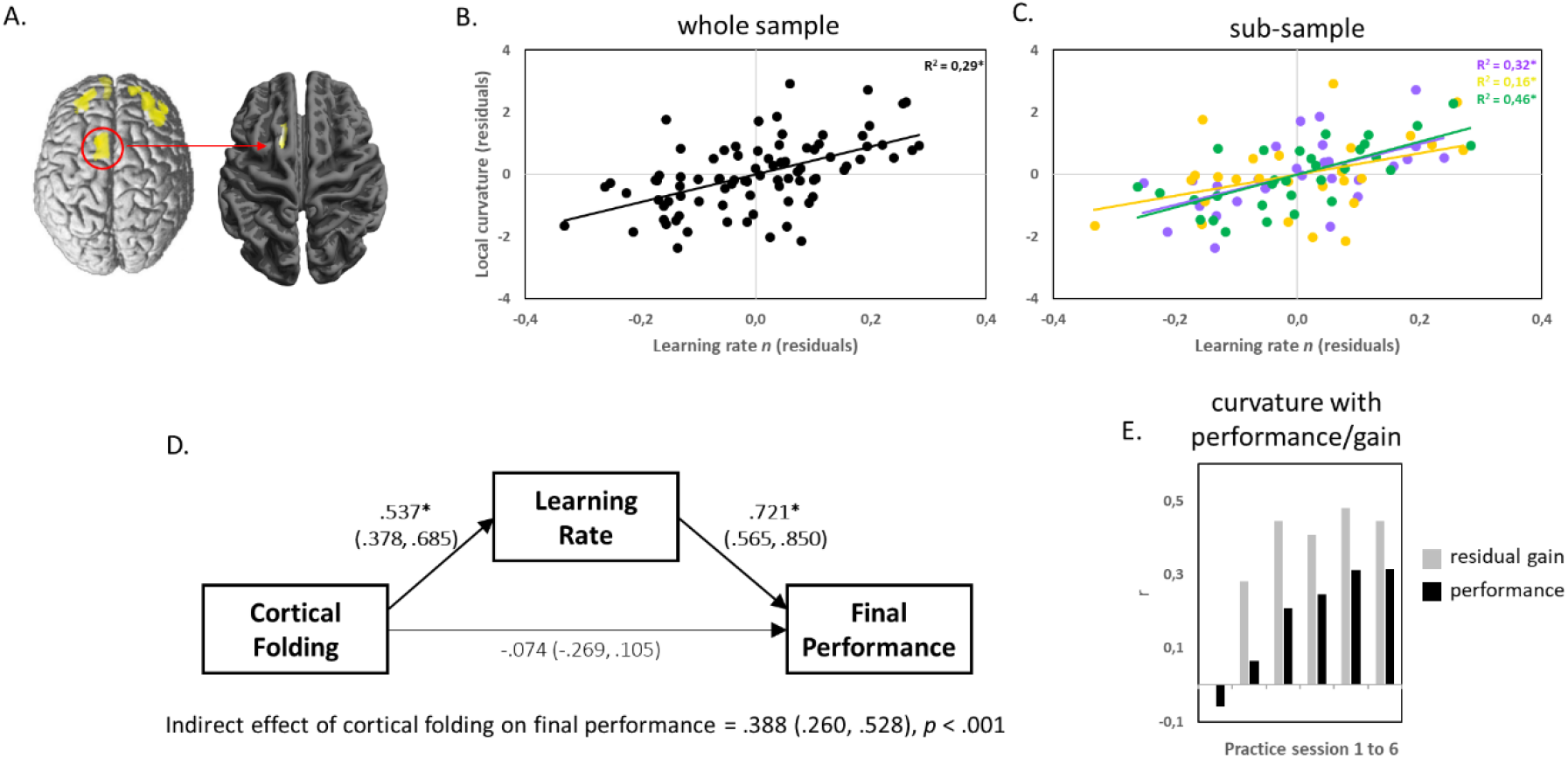
Cortical folding predicts learning in regions undergoing practice-induced structural plasticity. Relationship between folding and plasticity in the premotor cortex. (A) Positive correlation of cortical curvature in pre-SMA/SMA (in the cluster of significant learning-induced gray matter changes in ^35^) and learning rate. Practice-induced plasticity is depicted across the whole brain on the left side and the overlapping effect of cortical folding on learning rate is shown on the right side (only pre-SMA/SMA was significant). (B and C) Whole-sample and sub-sample correlations between learning rate and cortical curvature in the pre-SMA/SMA cluster in A. (D) SEM depicting the relationship between cortical folding in pre-SMA/SMA, learning rate (adjusted for *a*) and final performance on session 6. Standardized coefficients with 95% bootstrapped confidence intervals (CI) are represented on paths. (E) Pearson correlation coefficients between residualized cortical folding and motor performance. Grey bars represent session-specific performance controlled for initial performance in session 1 (i.e., residual gain) and black bars represent correlations with actual session-specific performance. * indicate significant correlations/paths at *p* < .05 (with CIs not including zero).

We averaged cortical curvature values within the previously identified pre-SMA/SMA cluster ^35^ (gray matter increase at MNI coordinate xyz -12 13 64, peak Z-value = 4.35). Average cortical curvature in this cluster predicted individual differences in learning rate (*R*^*2*^ = 0.29, *p* < .001 for the whole sample, Fig. 5B). This effect was consistent across the three sub-samples (Fig. 5C). Using SEM of ‘plasticity’ ROI values confirmed the effect of cortical folding on final performance (both unadjusted for *a*, Fig. 5D) that was mediated via learning rate *n* (adjusted for *a*).

### Morphology of tertiary sulci predicts learning rate

Recent studies have linked higher cognitive performance to the presence and prominence of tertiary sulci in the frontal cortex of human participants ^8,41,44^. Tertiary sulci are small, evolutionarily new cortical structures with great potential for identifying new connections between neuroanatomical substrates and human-specific aspects of cognition ^5^. Here we tested whether variations in the presence of tertiary sulci (PaM) in a region encompassing the left caudal superior frontal gyrus affect learning rate via its prominence (sulcal surface area) and folding characteristics.

We used the nomenclature and sulcus labeling methodology of ^58^ to manually define the sulcal landscape in the left caudal superior frontal gyrus (SFG) and adjacent precentral regions. We labeled 458 sulci in the left hemisphere (labeled sulci for each individual are shown in Figs. S12-S14). According to Germann et al. ^58^, the caudal SFG includes major sulci found in each individual brain (Fig. 6A): the interhemispheric fissure (IF), the superior precentral sulcus (SPr), the superior frontal sulcus (SFS), and the central sulcus (CS). The SPr in ^58^ appeared continuous for 76% of participants and was split in two branches for the remaining 24% of participants. One branch is usually caudal to the SFG and forms the base of the superior frontal sulcus, and the other branch is caudal to the dorsal portion of the middle frontal gyrus. Largely consistent with ^58^, we show a continuous SPr in the majority of participants in our sample (65%, 55 out of 84) and the two-branch pattern in 35% of participants (29 out of 84). Germann et al. ^58^ noted several smaller tertiary sulci that were heterogeneous in presence, appearance, and number (see also Fig. 6A): the medial precentral sulcus (MeP), the marginal precentral sulcus (MaP) as well as the paramidline sulci (PaM) with one or more short portions within caudal SFG oriented parallel to the SFS ^13,58^. While MeP and PaM were found in almost every participant within the boundaries of our region-of-interest in caudal SFG, MaP occurred in 66% of participants in ^58^ and in 50% of the participants (42 out of 84) in our sample. The cluster of vertices representing the effect of cortical curvature on learning rate *n* (see Fig. 2A) was anterior to MeP and medial to SFS (see white outline in one participant’s left hemisphere in Fig. 6A) and likely colocalized with PaM sulci. Thus, we tested the influence of PaM number and PaM morphology (folding index and surface area) on learning rate *n* using a structural equation model (SEM) that extends our above model (illustrated in Figure 3A).

**Fig. 6.**
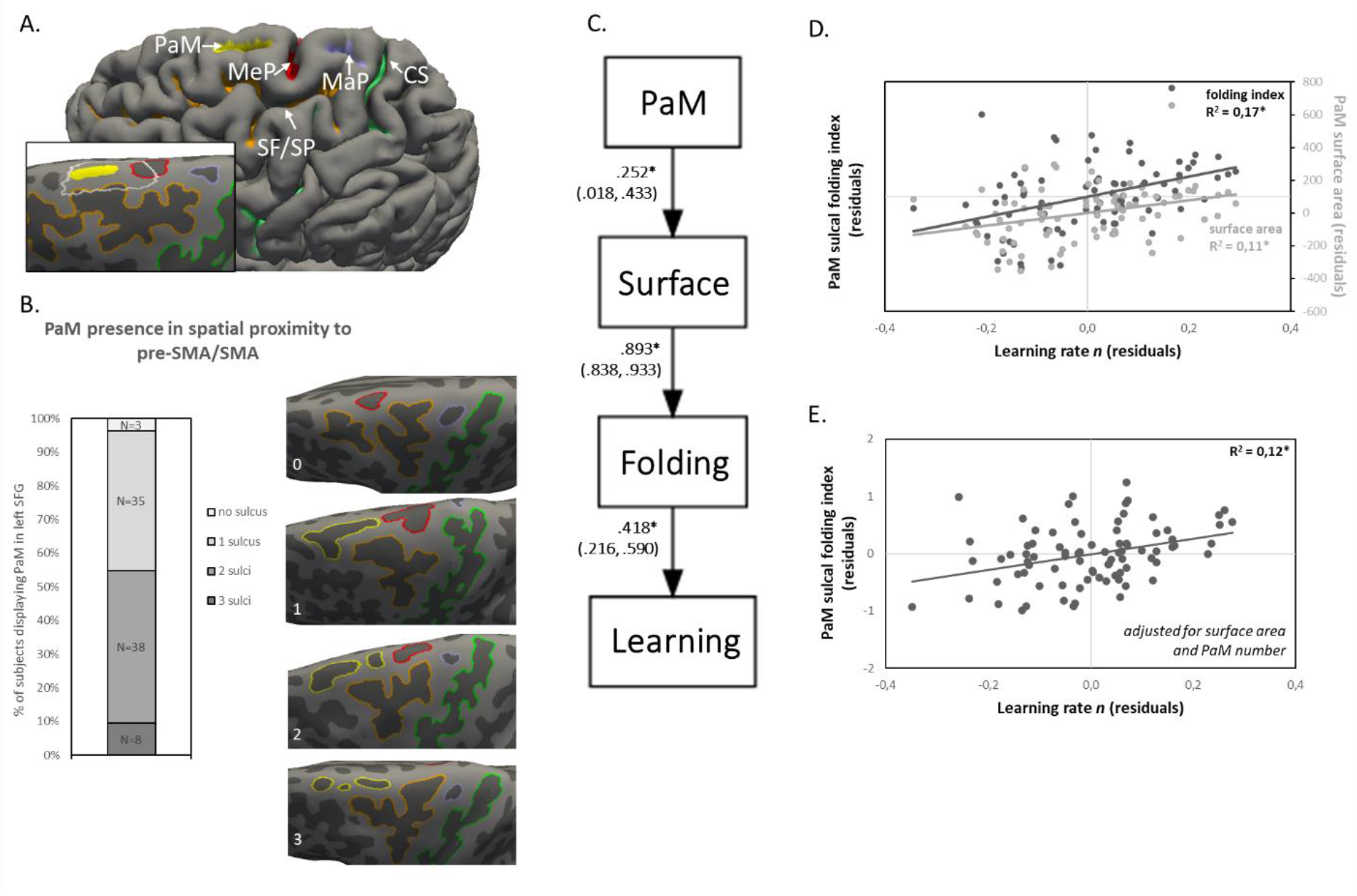
The morphology of tertiary sulci predicts individual learning rate. Analysis of tertiary sulci in the premotor cortex. (A) Labeling of cortical sulci in the left caudal superior frontal gyrus and adjacent precentral regions in a representative participant. Labeled left hemisphere sulci from all participants (n = 84) are shown in Figs. S12-S14. (B) The number of paramidline sulci for all participants is shown. The presence of PaM in the left caudal SFG (overlapping with the target cluster thresholded at p < .001) varies from PaM absence (no overlap between target cluster and PaM) in three participants to 3 PaM sulci in eight participants (Nabsence = 3, N1xPaM = 35, N2xPaM = 38, N3xPaM = 8). Surfaces of representative participants per number category (PaM are marked in yellow) are shown on the right-hand side. (C) A model representing the relationships between PaM sulcal folding index (“Folding”), PaM sulcal surface area (“Surface”), PaM sulcal number (“PaM”), and learning rate *n* (“Learning”). Standardized coefficients with 95% bootstrapped CIs are represented on paths. (D and E) Correlations between PaM sulcal folding index and learning rate. Folding index is either adjusted (D) or unadjusted (C) for differences in surface area and PaM presence. Note that all variables in the model and correlation analyses were corrected for differences in age, gender, height, head coil, study, baseline performance, and total intracranial volume. * indicate significant paths/correlations at *p* < .05 (with CIs not including zero).

PaM sulci number, PaM surface area, and PaM folding index were submitted to SEM to predict learning rate *n* (Fig. 6C). Significant relationships were found for (a) the presence of PaM (number) and PaM surface area, (b) PaM surface area and PaM sulcal folding as well as (c) PaM sulcal folding and learning rate *n* (Fig. 6C,D). Importantly, PaM number and surface area indirectly affected learning rate *n* via folding (presence: indirect effect of PaM number on *n:* 0.09 [95% CI = .006, .192], *p* = .046; prominence: indirect effect of PaM surface area on *n:* 0.37 [95% CI = .186, .538], *p* < .001). Moreover, a partial correlation analysis revealed a significant positive correlation between PaM sulcal folding and *n* when controlling for PaM number and PaM surface area (partial *R*^*2*^ = 0.12, *p* = .001, Fig. 6E).

## Discussion

Given the complexity of mechanisms involved in the expansion and folding of the cerebral cortex, and thus its tremendous costs in terms of genetic, cellular, and histogenic evolution, the ecological advantages of cortical folding must be more than remarkable ^59^. Using longitudinal training data, we show that human participants with higher degrees of cortical folding in premotor areas have larger performance gains (steepness of the learning rate) across several sessions of motor practice. Cortical folding had an indirect effect on attained performance levels via its strong impact on performance gain. The observed local associations between performance gain and cortical folding overlapped with practice-induced structural plasticity in premotor areas and with the morphological characteristics of hominoid-specific tertiary sulci. Higher cortical folding was related to larger cortical surface area, but not at the expense of lower cortical thickness or intracortical microstructure. Our results support the hypothesis that higher levels of cortical folding endow individuals with enhanced adaptive capacities, but not with superior performance per se.

Interindividual differences in global and local folding metrics were correlated with behavioural performance scores in previous studies involving adult humans ^8,19,24,37,38,43,44,60–63^. These studies usually assessed cognitive or memory performance at a single point in time – with intelligence quotient being the most commonly assessed variable to date. The effect size of brain-behaviour correlations varied considerably but generally suggest a positive association between higher folding and performance. Using a longitudinal measure of performance change, we report that approx. 30% of variance in learning rate are predicted by the degree of local cortical folding in premotor cortical regions (pre-supplementary/supplementary motor areas). In line with ^24^, larger cortical surface area contributed to the folding effect on learning but there was an additional significant surface area-independent contribution to cortical folding’s relationship with learning (Figs. 3 and 6). While the technical reproducibility of the folding-learning relationship (Fig. S6) was expected because of the high stability of non-invasive markers of external brain morphology, we were surprised by the consistency of positive correlations within an independent region-of-interest and for smaller sub-samples (Fig. 5C and S7).

We report a comparably large effect size for a brain behavioural study of cortical folding in adult humans (see correlation coefficients represented in both ROI and vertex-wise analyses, Figs. 2 and S5). The comparatively long practice time could have favored the identification of brain-behavioural relationships^64^. We found that associations between cortical folding and motor performance increased with practice. This can be explained by the increasing impact of residual gains on absolute performance levels across practice (Figs. 2E, 5E, S6D). In fact, performance gain mediated the effect of cortical folding on final performance (Figs. 2D and 5D). This suggests that cortical folding effects on acquired performance level may be an indirect consequence of cortical folding’s relationship with an underlying learning ability.

Practice augments individual performance differences which are associated with relatively stable factors (e.g., aptitude, genotype, phenotypic, and other psychological traits), a view held in developmental psychology and behavioural genetics ^32,65^. We interpret our finding as a reflection of interindividual differences in capabilities (rather than actual performance levels), mediated by the degree of cortical folding ^28,29^. Although in prospective cohort studies, within-person change trajectories generally have lower heritability rates than cross-sectional measurements obtained from different groups of individuals ^66^, the stimulus for performance change is usually under experimental control in an intervention study. Intervention studies from behavioural genetics report higher heritability rates with motor practice ^31,32^. In particular, the twin study by Williams & Gross ^31^ used the same postural learning task as in the present work (stabilometer task) and found increased genetic influences on motor performance across six practice sessions. When our participants learned the same postural task (also across six sessions), the impact of initial performance differences on subsequent achievements decreased during practice (Fig. S15). A large portion of this increasing residual variance during practice was explained by variations in cortical folding of the pre-SMA/SMA. Future studies are required to disentangle the specific contributions of genetic or (early) environmental factors to behaviourally meaningful variations in cortical folding.

A large network of cortical and sub-cortical regions is involved in gait and postural control ^67^ and our analyses specifically focused on the cerebral cortical contributions to individual differences in postural learning. The supplementary motor area is critically involved in anticipatory postural control and gait ^68,69^. This region also adapts its structure in response to postural training ^70^. Practice of the stabilometer task (used in the present work) induces structural gray matter changes in the left pre-SMA/SMA and microstructural changes in the underlying white matter tracts of the left centrum semiovale^35^. Practice-induced structural changes were also accompanied by increased functional connectivity between the pre-SMA/SMA and medial parietal areas ^71^. This indicates that postural learning is associated with the connectivity and folding pattern of the pre-SMA/SMA embedded within a wider cortical-subcortical network responsible for posture and gait control.

An individual’s genotype has a significant impact on practice-induced motor performance gains ^32^ as well as on sulcal morphology ^39^. To which extent variations in cortical folding are predictive of learning success in different types of motor tasks remains unclear. The folding-learning associations observed in our study suggest a comparably homogeneous effect within some selected anthropometric, demographic, and performance sub-categories of our sample (Fig. S8). Although the overall pattern of cortical folding is relatively stable across life, supportive interventions could have a significant impact on motor learning. In line with this, we found an overlap of meaningful folding variations with practice-induced plasticity in pre-SMA/SMA which is consistent with research using juggling as long-term motor learning paradigm^25^. A spatial overlap was found between juggling-induced gray matter changes in parietal regions and an association between baseline parietal gray matter volume with subsequent learning-induced performance improvements^25^. Together, this supports future efforts to mitigate potential behavioural deficits related to cortical predispositions by using appropriate training methods. Second, additional interventions such as vigorous physical exercise in the weeks prior to motor practice can further improve learning in this particular postural task ^47^. The beneficial effect of vigorous exercise on postural learning is mediated by structural and functional changes in the fronto-parietal brain network ^47,72^. Thus, plasticity-inducing intervention strategies may be a fruitful approach to enhance learning beyond neural predispositions (see Supplementary text).

Cortical folding is the result of different mechanisms extrinsic and intrinsic to the cortical sheet. Extrinsic sources can be the volumetric constraints of the cranial vault harboring an expanded cortex or connected axons pulling cortical and sub-cortical regions closer together to enhance information transmission speed ^73^. Intrinsic mechanisms can be a higher level of cortical neurogenesis, differential tangential expansion of upper cortical layers or neuropile growth ^4,6^. Cross-species comparisons show that humans possess a remarkably large number of neurons in the cerebral cortex ^74^. Studies in ferret, macaque and human brain found that, in species with a folded cortex, the rate of neurogenesis is heterogeneous along the developing cortical mantle ^4^. Higher rates of neurogenesis emerging in upper cortical layers of human-specific gene knock-in mice ^18^ result in cortical buckling of the otherwise lissencephalic mouse brain and in better spatial learning capabilities in these animals. In addition, neuropile expansion influences the growth of late developing cortical regions (e.g. tertiary sulci) ^6^. Thus, higher adaptive requirements of the postural system during development or evolution could have fostered surface expansion and folding in task-specific cortical regions ^1,75–77^. Our study revealed that intra-specific variations in cortical folding and tertiary sulcus morphology predict learning of a challenging postural task. The results also show that the impact of cortical folding on learning is related to differences in cortical surface area as well as surface area-independent extrinsic and/or intrinsic factors of folding, but not to differences in intracortical microstructure (Fig. 3, 4 and 6). The pattern of correlations in Fig. 3 and 6 indicates that (1) sulcal and gyral surface area exerts both significant positive and negative influences on learning rate, (2) the significant positive influence of surface area on learning rate is mediated via its impact on cortical folding and (3) the significant negative influence of surface area on learning rate likely arises from the gyral regions around the tertiary paramidline sulci. Further studies with higher-resolution MRI techniques are required to disentangle the contributions of extrinsic and intrinsic sources of cortical folding (e.g. U-fibres, layer-specific microstructure) and of gyral, sulcal and fundal points^78^ to behavioural differences.

While the underlying factors of cortical folding are subject to intense research in the biological and physical sciences ^79^, our study investigated the behavioural capacities that are enabled by higher levels of local cortical folding in humans. Cortical folding was related to learning rates over multiple motor practice sessions. The fact that the learning rates were adjusted for differences in initial performance (and that cortical folding was also not related to initial performance differences) has implications for inclusive learning approaches. Individual learning capabilities, irrespective of initial performance conditions, may be associated with stable and region-specific morphological characteristics of the cortex. Under the assumption of physical constraints to the information processing capacity of the cerebral cortex ^9^, education seems critical for an individual to realize its potential in a particular domain regardless of their initial performance in that domain. Our study also showed that learning rates mediated between higher cortical folding and asymptotic levels of performance at the end of a practice period. In that sense, improved human performance does not necessarily emerge from an extraordinary brain morphology, but rather from an interaction between fertile learning environments and remarkably high learning capabilities ^29^. In our study with healthy human participants, high learning capability was partially reflected in the surface morphology of the human neocortex.

## Methods

### Experimental Design

We analyzed MRI and behavioural data from three independent motor learning experiments involving adult human participants (see Participants). All participants with complete MRI and behavioural data from these three studies were included in the analyses. MRI of the brain was performed before motor practice of a challenging new postural task on a stabilometer (see Postural task practice). Indices of motor performance and learning rate over several practice sessions (see Analysis of motor learning) were correlated with local indices of cortical folding from preprocessed MRI data (see MRI acquisition and MRI preprocessing). Statistical analyses involved vertex-wise comparisons of cortical curvature and region-of-interest (ROI) comparisons of cortical and sulcal morphology as well as intracortical microstructure (see Statistical analysis).

### Participants

A sample of 131 right-handed participants with normal or corrected-to-normal vision (mean age of 24.6 years, age range of 19-35 years, 57 females, mean body height 174 cm, body height range 153-191 cm) was included from the datasets of three independent motor learning experiments ^35,46–48^. In addition, data from ^80^ was used to increase the sample size for the analysis of short-term improvements in motor performance (only data for session 1). The studies were performed in accordance with the Declaration of Helsinki and approved by the Ethics Committees of the Universities of Leipzig and Magdeburg (Germany). Exclusion criteria were contraindications to magnetic resonance imaging (MRI), body mass index (BMI) > 30 kg/cm^2^, a history of neuropsychiatric diseases, left-handedness and prior experience with the task to be learnt. Participants were screened for contraindications of MRI before participation. Participants were naive to the experimental setup and postural training procedure and were of comparable educational level (all participants had A-level).

### Postural task practice

Participants learned a challenging whole-body postural task on a stabilometer either on one practice session (N=131) or over six practice sessions (N=84). From the 84 participants, practice sessions were either distributed over six consecutive weeks with one training session per week (N=58, study 1 and study 3) or distributed over four consecutive weeks with 1-2 practice sessions per week (N=26, study 2). The stabilometer is a movable, seasaw-like platform attached to a superimposed pivot with a maximum board deviation of 26° to each tilt side (stability platform, model 16030L, Lafayette Instrument). Participants were instructed to stand on the stabilometer board and hold/restabilize the platform within a tolerance interval of +-3° from the horizontal (see Supplementary Video files). After each of the 15 trials (30 seconds in each trial) per practice session, participants received verbal performance feedback. Performance was measured as accumulated time (in seconds) participants were able to maintain the platform in the +-3° tolerance interval (Time-in-balance). A short break of 2 minutes between trials was used to avoid fatigue. Each practice session lasted approx. 45 minutes. To familiarize subjects with the task and to prevent falls, we allowed the use of a supporting hand rail in the first trial of session 1. Familiarization trials were excluded from the analysis. We used a discovery learning approach ^81^ in which no information about the performance strategy (only the trial-wise quantitative performance feedback) was provided during practice. Therefore, participants had to discover their optimal strategy to improve task performance (e.g. error correction strategy with legs, hip, and arms) based on by trial and error.

### Analysis of motor learning

The mean performance scores (mean of time-in-balance values across 15 trials) on each of the six practice sessions for each individual participant were fitted to a general power function, *y(x) = a * x*^*n*^, which describes motor learning over longer timescales well ^82^. In this function, the base *a* denotes initial task performance, *x* is training session (time devoted to practice), and the exponent *n* indicates the slope of the function (rate of learning). Furthermore, early learning was calculated from performance data on the first practice session. For that, we subtracted the mean of the first five trials from the mean of the last five trials. We used learning rate (*n*), initial performance (*a*) and early learning (performance gain during session 1) as dependent variables in statistical analyses of brain-behavioural relationships. As expected from motor learning literature ^50^, initial performance negatively predicted learning rate (Fig. S1). To get an unbiased readout of learning ability, we adjusted *n* for differences in *a* ^*51*^.

### Magnetic resonance imaging (MRI) acquisition

Anatomical T1-weighted MPRAGE data ^83^ were acquired on a 3T MAGNETOM magnetic resonance imaging (MRI) system (Siemens Healthcare) with 176 slices in sagittal orientation (study 1 N=27: Tim Trio system using a 32-channel head coil, study 2 N=26: Prisma system using a 64-channel head coil, study 3 N=31: Prisma system using a 32-channel head coil). The imaging parameters used were as follows. Study 3: inversion time (TI) = 900 ms, repetition time (TR) = 2300 ms, echo time (TE) = 2.98 ms, flip angle = 9°, field-of-view (FOV) = 256 x 240 mm^2^, spatial resolution = 1 x 1 x 1 mm^3^; study 1: (TI) = 650 ms, (TR) = 1300 ms, (TE) = 3.46 ms, flip angle = 10°, (FOV) = 256 x 240 mm^2^, spatial resolution = 1 x 1 x 1 mm^3^; study 2: (TR) = 2600 ms; (TE) = 5.18 ms; flip angle = 7°; (FOV) = 256 x 256 mm^2^; spatial resolution = 0.8 x 0.8 x 0.8 mm^3^. Due to the potential influence of the radiofrequency head coil on brain morphometric indices ^84^ we corrected for this factor in the statistical models. In addition, we corrected for MRI scanner and MPRAGE sequence-specific effects using a separate nuisance covariate for each of the three studies.

### MRI preprocessing

MR images of all participants passed both the visual quality inspection and the CAT12 data quality checks. All scans from 131 participants reached a weighted average image quality rating (IQR) of 86.79% (range 80.64%–89.87%) corresponding to a quality grade B while the long-term practice cohort (N=84) reached a weighted average (IQR) of 87.32% (quality grade B; range 85.62%-89.87%). T1-weighted images were preprocessed using the CAT12 toolbox, v12.7 r1738 (Christian Gaser, Structural Brain Mapping Group, Jena University Hospital; http://www.neuro.uni-jena.de/cat12/, ^85^) within SPM12 v7771 (Statistical Parametric Mapping, Wellcome Trust Centre for Neuroimaging; http://www.fil.ion.ucl.ac.uk/spm/software/spm12/) for Matlab R2017b (The MathWorks, Inc.). This image analysis pipeline allows for the computation of surface-based parameters based on, e.g., the mean curvature and procedures are described in detail on the CAT 12 website and manual (https://neuro-jena.github.io/cat/index.html#DOWNLOAD). All procedures followed the recommendations in the CAT 12 manual. Briefly, initial voxel-based processing involves spatially adaptive denoising, resampling, bias correction, affine registration and unified segmentation and provides starting estimates for subsequently refined image processing. Output images were then skull-stripped, parcellated into left and right hemisphere, cerebellum and subcortical areas as well as corrected for local intensity differences and adaptively segmented followed by spatial normalization. Subsequently, central cortical surfaces were reconstructed and topological defects were repaired using spherical harmonics. The refined central surface mesh provided the basis for extraction of local cortical folding metrics (e.g., local curvature) and resulting local values were projected onto each mesh node. Local gyrification ^52^ is revealed through estimations of “smoothed absolute mean curvature” based on averaging curvature values from each vertex of the surface mesh. Mean curvature is an extrinsic surface measure and represents change in direction of surface normals along the surface (normal are vectors pointing outwards perpendicular to the surface). Large negative values correspond to sulci and large positive values to gyri. The resulting values were averaged within a distance of 3 mm and converted to absolute values (both sulcal and gyral regions have positive values, see ^52^). We then applied a surface-based heat kernel filter with FWHM = 20 mm, as recommended for vertex-wise gyrification in the CAT12 user manual. The resulting values give information about the local amount of gyrification. Finally, individual central surfaces were registered to the Freesurfer “FsAverage” template using spherical mapping with minimal distortions. Local gyrification values are transferred onto this FsAverage template.

To assess local interactions of cortical folding, surface area and cortical thickness in the left caudal superior frontal gyrus and to manually define and label sulci in individual subjects native space, we additionally used FreeSurfer automated segmentation tools ^86,87^ (FreeSurfer 6) to reconstruct cortical surfaces (recon-all command; https://freesurfer.net/fswiki/recon-all) from all baseline T1-weighted MRI images of the long-term practice cohort (N=84). Cortical reconstruction and volumetric segmentation were performed with the Freesurfer image analysis suite, which is documented and freely available for download online (http://surfer.nmr.mgh.harvard.edu/). The technical details of these procedures are described on the FreeSurfer website (https://surfer.nmr.mgh.harvard.edu/fswiki/FreeSurferMethodsCitation). Briefly, this processing includes motion correction of volumetric T1 weighted images, removal of non-brain tissue using a hybrid watershed/surface deformation procedure, automated Talairach transformation, segmentation of the subcortical white matter and deep gray matter volumetric structures (including hippocampus, amygdala, caudate, putamen, ventricles) intensity normalization, tessellation of the grey matter white matter boundary, automated topology correction, and surface deformation following intensity gradients to optimally place the grey/white and grey/cerebrospinal fluid borders at the location where the greatest shift in intensity defines the transition to the other tissue class. Once the cortical models are complete, a number of deformable procedures can be performed for further data processing and analysis including surface inflation, registration to a spherical atlas which is based on individual cortical folding patterns to match cortical geometry across subjects, parcellation of the cerebral cortex into units with respect to gyral and sulcal structure, and creation of a variety of surface-based data including maps of curvature and surface area. This method uses both intensity and continuity information from the entire three-dimensional MR volume in segmentation and deformation procedures to produce representations of cortical thickness, calculated as the closest distance from the grey/white boundary to the grey/CSF boundary at each vertex on the tessellated surface. The maps are created using spatial intensity gradients across tissue classes and are therefore not simply reliant on absolute signal intensity. The maps produced are not restricted to the voxel resolution of the original data thus are capable of detecting submillimeter differences between groups. Procedures for the measurement of cortical thickness have been validated against histological analysis and manual measurements. We supplemented the analysis of local cortical geometry (curvature) with an analysis of a gyrification metric that depends on the ratio between the outer hull surface area and the local cortical surface area (called outer-surface-based gyrification indices). Therefore, we computed the local gyrification index ^40^ of freesurfer cortical reconstructions.

Based on the group-level result of a correlation between motor learning ability and local cortical curvature in the left pre-SMA/SMA (Fig. 2A), we manually defined a region-of-interest (ROI) in the left caudal SFG (including pre-SMA/SMA) encompassing the cortex in SFG extending from the anterior edge of the superior precentral sulcus (joining the medial precentral sulcus) to the caudal part of the superior frontal sulcus (at the level of the gyral bridge between middle and superior frontal gyrus) and, in the medio-lateral dimension, the cortex running from the interhemispheric fissure to the superior frontal sulcus ^58^ on the Freesurfer “FsAverage” template brain. This ROI was projected to each participant’s native space and local indices of cortical folding ^88^, cortical surface area and cortical thickness were extracted from the white matter surface (to avoid blood vessel contamination ^8^) and averaged in this ROI. In addition to that, we manually defined the sulcal landscape in the left caudal SFG using the freeview tool in FreeSurfer and the labeling methodology of ^58^. The following sulci were investigated: the superior precentral sulcus (SP), the superior frontal sulcus (SF), the central sulcus (CS), the medial precentral sulcus (MeP), the marginal precentral sulcus (MaP) and the paramidline sulci (PaM). Based on ^58^, we first drew the sulcal lines on the inflated cortical surfaces and validated the position and shape of each sulcus using the corresponding pial surface image ^8^. Thus, information from the inflated and pial surfaces informed our labeling and allowed us to form a consensus across surfaces and clearly determine each sulcal boundary. Although our analysis focused on PaM, we manually identified all sulci in the caudal and superior part of the lateral frontal cortex (in total 458 sulci in left hemispheres; labeled sulci from each individual are depicted in Figs. S12-S14) to ensure the most accurate definition of PaM components ^13,58,89^.

The superior frontal gyrus of the human brain typically contains three PaM components (anterior, intermediate and posterior component) that are arranged in parallel or orthogonal to and in-between the interhemispheric fissure and the superior frontal sulcus ^13,58,89^. We focused our analysis on the posterior and intermediate PaM components that are located in close spatial relationship to the pre-SMA/SMA. PaM sulci were located on the lateral surface of the left hemisphere, medial to SF, anterior to SP and MeP. We labeled PaM sulci which overlap with the cluster found in the group-level analysis (Fig. 2; group-level cluster was projected to individual surfaces).

Next, we quantified the surface area and folding index of each labeled PaM sulcus using mris_anatomical_stats function included in FreeSurfer. In case of more than one identified PaM sulcus per hemisphere, we added surface area and folding values.

### Statistical analysis

Our main goals were to test for positive relationships between inter-individual differences in learning rate or motor performance with local cortical folding. In these analyses, we corrected for the influence of age, gender, body size, total intracranial volume (estimated using CAT12 module “Estimating TIV”) and study (initial differences in *a* were only adjusted in the analysis of learning rate).

#### Motor behaviour

Short-term changes in motor performance (time-in-balance in seconds) in the first practice session (*N*=131) were analyzed with repeated measures analysis of variance (RM-ANOVA) with within-subject factor TRIAL (15 levels) in SPSS (IBM SPSS Statistics, Version 28.0.1.0, Armonk, NY). Long-term changes in motor performance across the six practice sessions were analyzed with RM-ANOVA of the session mean values (mean of 15 trails per session) with within-subject factor SESSION (6 levels). Trial-to-trial variation in performance were calculated with the coefficient-of-variation (COV, standard deviation divided by the mean) for each session and subjected to RM-ANOVA with factor SESSION (6 levels). Session-specific inter-individual variation was quantified using interquartile range between the upper and lower 25% of mean performance values. Pearson correlations were used to relate mean performance values across sessions.

#### Analysis of cortical folding on long-term learning, initial performance and short-term adaptation

Our main predictions were tested with a multiple linear regression model in SPM12 (http://www.fil.ion.ucl.ac.uk/spm/software/spm12/) with local cortical folding values across the cortex as dependent variable and learning rate *n* (*N*=84, corrected for individual differences in initial performance level *a*) or initial performance *a* as well as short-term adaptation (*N*=84 and *N*=131) as predictors. In each analysis, we corrected for the influence of age ^90^, gender ^91^, body height ^46^, head coil ^84^, total intracranial volume ^92^ and training study ^35,47,48^. Covariation between (nuisance) variables are shown in Fig. S2. Statistical inference of positive relationships between behavioural parameters and cortical curvature was performed across the whole cortex (exploratory analysis) with non-parametric permutation test (vertex-level T-statistics) and 5000 permutations. *p*-values were considered significant at an FWE corrected threshold of p < 0.05. Technical reproduction of significant effects was performed using a further MRI scan from the same participants. This further MRI scan was obtained after the last motor practice session either six weeks (study 1 and study 3) or four weeks (study 2) after the baseline MRI scan. The cluster extent from the initial exploratory whole-cortex analysis (Fig. 2A) was used as inclusive mask and surface measures from the second time point were averaged in this respective mask. Cortical folding values in this mask were highly reliable across the two MRI time points (*r* = 0.964). The overlap between cortical folding and practice-induced plasticity in grey matter volume was calculated using a group-space mask of the cluster in pre-SMA/SMA where we previously identified grey matter changes across the six-week practice period ^35^ (xyz MNI coordinate -12, 13, 64, cluster with highest Z-value=4.35 across the whole brain). The voxel-space cluster (rendered brain see Fig. 2) was projected to the FsAverage surface template using CAT12 surface tools. The cortical folding values in this mask as well as in the mask for technical replication were averaged and subjected to statistical analysis in SPSS. In subsequent correlation analyses, we used residualized learning rate and cortical curvature values (corrected for age, gender, initial performance, body height, head coil, TIV, training study) to determine reproducibility, effect sizes and coincidence of folding and plasticity (Figs. 2, 5 and S6). Variations of the effect were tested with Pearson correlation analyses of the positive relationship between learning rate and cortical folding in pre-SMA/SMA (time point 1) in differently categorized sub-groups of the original sample (N=84, Fig. S8). We categorized this sample with respect to the following demographic, anthropometric, sub-group-related, activity-related and performance-related variables by means of binarized dummy variables or median split: age, gender, body height, initial performance level, physical activity level (above or below 4 hours per week), vigorous physical exercise in 2 weeks prior to motor practice, study-specific sub-groups. For each correlation of the categorized groups, we used residualized learning rate and cortical folding variables with the categorization variable not included in the residualization procedure.

In addition to the main cohort (N=84) we included additional 47 participants from ^46^ in the correlation of initial performance and short-term adaptation with cortical folding. These additional participants were measured on a Tim Trio MRI system using either 12-channel or 32-channel head coil (which was corrected for in the respective statistical model, for more details see ^46^). Dependent variables were either initial performance (mean performance of 15 trials in practice session 1) or early learning calculated as the difference between the mean of the last 5 trials and the mean of the first 5 trails from practice session one.

#### Myelin-sensitive magnetization transfer saturation (MT) and estimates of neurite density index (NDI) from neurite-orientation-and-dispersion-imaging (NODDI) modeling of diffusion MRI

Myelin-sensitive MT values were calculated from multiparametric quantitative MRI protocol with 0.8 mm isotropic voxel size ^48^ and NDI values were calculated from NODDI modeling of diffusion MRI data with 1.6 mm resolution ^93^ within the gray matter in the study2-subsample (N=26). Both MT and NODDI metrics are highly reliable ^48,93^ and calculation of NDI values within gray matter was adjusted according to ^94^. Based on previous findings ^1^, MT values were extracted and averaged within three cortical depth-dependent tissue compartments (superficial and deep cortical gray matter [GM] and cortex-adjacent white matter) in individual space using CAT12 surface tools. For each compartment, a mean sampling function (average along surface normal) and a equi-distance mapping model with 7 steps was employed (startpoint: superficial=-0.5, deep=0, white matter=0.5; endpoint: superficial=0, deep=0.5, white matter=1.0). Superficial GM extends from the gray matter/CSF border to the central surface. Deep GM extends from the central surface to the gray/white matter border and the cortex-adjacent white matter extends from the gray/white matter border into the cortex-adjacent white matter. Due to lower resolution of diffusion data, NDI values were sampled from the whole GM compartment (startpoint=-0.5, endpoint=0.5). Resulting MT maps and NDI maps were resampled into template space and smoothed with filter size of 15 mm FWHM. To visualize MT/NDI distribution across the whole cortex (Fig. 4A,B), we additionally mapped and averaged MT values in the whole gray matter compartment (from gray matter/CSF to gray/white matter boundary). For statistical analysis, compartment-specific values were extracted from the region overlapping with the pre-SMA/SMA cluster (Fig. 2A), but also analyzed vertex-wise. We used residualized (corrected for age, gender, body height, TIV, initial performance) MT/NDI, learning rate, cortical folding and age parameters for all Pearson and partial correlation analyses or adjusted for these nuisance variables in SPM statistical models for vertex-wise analyses (except of the age by MT/NDI correlation in which we did not correct for age and initial performance).

#### Structural equation modeling (SEM)

SEM was used to better understand the dependencies between motor behaviour and cortical folding (Figs. 2 and 5) as well as between cortex morphology variables (Fig. 3 and 6). For this purpose, we used the lavaan package ^95^ running in R (i386 4.1.1, R Core Team, 2020) and RStudio. In the first model (Fig. 2D), cortical folding in the pre-SMA/SMA and residualized learning rate *n* were used as exogenous variables to predict final performance in practice session 6 (SEM fit indices *RMSEA* = 0.000, *SRMR* = 0.000, *CFI* = 1.000, *TLI* = 1.000). Note that values of cortical folding and final performance were not adjust for differences in initial performance in this analysis. In the second model (SEM fit indices *RMSEA* = 0.000, *SRMR* = 0.000, *CFI* = 1.000, *TLI* = 1.000, Fig. 5) we used the independent ROI in which practice-induced gray matter changes were found previously ^35^ (Fig. 2G-I). In the third model (Fig. 3, SEM fit indices *RMSEA* = 0.000, *SRMR* = 0.000, *CFI* = 1.000, *TLI* = 1.000), surface area, cortical thickness and cortical folding indices in the left caudal SFG were used as exogenous variables to predict learning rate *n*. In the fourth model (Fig. 6, SEM fit indices *RMSEA* = 0.000, *SRMR* = 0.019, *CFI* = 1.000, *TLI* = 1.025), the number of PaM sulci, PaM surface area and PaM folding index were used as exogenous variables to predict learning rate *n*. All values were residualized for age, gender, body height, TIV, initial performance and training study with the exception that values of cortical folding and final performance in the first two models were not adjust for differences in initial performance. We calculated direct and indirect effects with 95% bootstrapped CIs using 5000 permutations.

## Funding

German Research Foundation grant SFB 1436/C01 (MT) German Research Foundation grant SFB 1436/C01 (GZ)

## Author contributions

Conceptualization: MT, NL

Methodology: MT, GZ

Investigation and formal analysis: MT

Visualization: MT

Data curation: MT

Writing—original draft: MT

Writing—review & editing: MT, GZ, NL

## Competing interests

Authors declare that they have no competing interests.

## Data availability

All data are available in the main text or the supplementary materials. In addition, data and code used for this project have been made freely available under https://doi.org/10.24352/UB.OVGU-2023-095. Visualizations of all sulcal definitions generated for each participant are provided in the Supplementary Materials. Requests for further information or raw data should be directed to the corresponding author, M.T.

## Notes

https://doi.org/10.24352/UB.OVGU-2023-095

### Competing Interest Statement

The authors have declared no competing interest.

